# Regulatable In Vivo Gene Expression via Adaptamers

**DOI:** 10.64898/2025.12.21.695777

**Authors:** Jack Bryant, Laura Herron, Yesh Doctor, Finn McRae, Jocelyn Wang, Theodore Hirsch, Andrew Portell, Satheesh Kumar, Prashant Mali

## Abstract

Precise, reversible control of transgene expression is essential for safe and durable gene therapy, yet current inducible systems remain difficult to translate in vivo due to large size, limited induction duration, and dependence on immunogenic regulators. Here we present the **adaptamer** (<u>AD</u>AR modulatable <u>aptamer</u>), a compact (<120 bp) RNA switch that couples an FDA-approved small-molecule–responsive aptamer with endogenous ADAR–mediated RNA editing to regulate expression. Ligand binding stabilizes a double-stranded RNA structure that recruits ADAR to convert a stop codon into a sense codon, restoring downstream protein translation. This cleavage-free, post-transcriptional mechanism enables precise, small-molecule–dependent modulation without exogenous protein machinery. We demonstrate that adaptamers are highly functional across multiple cell lines, including human T-cells. In mice, AAV-delivered adaptamer-controlled FGF21 expression induced metabolic remodeling, significantly increasing energy expenditure and reversing obesity. This minimal, programmable system offers a clinically compatible approach for tunable genetic medicines and safe deployment of pleiotropic and dose-limited proteins.

## INTRODUCTION

Gene and cell therapies have emerged as powerful modalities for treating a wide range of diseases; however, they generally lack precise control over transgene expression, restricting their use to transient expression, irreversible “hit-and-run” genomic modifications, or permanent constitutive expression (*1–4*). Because many therapeutic proteins are beneficial only within narrow dose and temporal windows—where both insufficient expression and excessive exposure can cause pathology—this limitation substantially narrows the spectrum of treatable diseases (*2*). Moreover, redosing of both viral and non-viral delivery systems remains impractical due to immunogenicity and cost. Consequently, optimal therapies for chronic diseases require vectors that persist long term while enabling tightly regulated, inducible expression (*2*).

Numerous protein-based genetic switches have been developed to meet these criteria, most notably the Tet-On system; however, their reliance on bacterial or engineered proteins can provoke immune responses that limit their utility in vivo (*5*, *6*). Riboswitches offer an alternative approach. These synthetic RNA elements, composed of a small-molecule-binding aptamer coupled to an effector domain, avoid adaptive immune activation because they are entirely RNA-based (*7–9*). In mammalian systems, however, riboswitches exclusively rely on small-molecule-regulated RNA cleavage or splicing to control gene expression (*7–9*). A cleavage based mechanism intrinsically limits the durability of induction, as the regulated RNA is rapidly degraded following small-molecule withdrawal, necessitating frequent dosing that may be unsafe or impractical in clinical settings (*10*). Splicing based mechanisms on the other hand rely on surrounding context to function properly making them transgene and vector dependent (*11*). In addition, highly engineered riboswitches—such as the recently reported pA regulator, which exceeds 500 base pairs—further constrain adeno-associated virus (AAV) packaging capacity and introduce complex RNA secondary structures that can impair translation and splicing efficiency (*9*). Dependence on cleavage or splicing also precludes deployment in RNA-based expression platforms: cleavage-based designs would self-destruct prior to delivery, while splicing-dependent switches remain inactive due to the absence of cytosolic splicing machinery. As a result, delivery approaches such as lentiviral vectors and mRNA lipid nanoparticles are excluded. Reflecting these challenges, only a single riboswitch, K19, has been evaluated in a mouse disease model; owing to rapid RNA turnover, therapeutic efficacy required twice-daily small-molecule dosing—a regimen likely to be intolerable in humans (*10*).

Adenosine deaminases acting on RNA (ADARs) have recently emerged as promising tools for programmable RNA editing (*12*, *13*). These enzymes are conserved across metazoans and catalyze the deamination of adenosine to inosine, which is interpreted as guanosine by the translational machinery. In humans, three catalytically active isoforms exist: the ubiquitously expressed, nuclear ADAR1 p110; the interferon-inducible, cytoplasmic ADAR1 p150; and the nervous system-enriched, nuclear ADAR2 (*14*). Although the rules governing ADAR targeting are still being elucidated, their strong preferences for specific RNA sequences and secondary structures are well characterized (*15*, *16*). This property has been exploited to conditionally silence engineered stop codons upstream of genes of interest, enabling expression in response to target transcripts or endogenous molecules (*17–20*). However, most such systems require exogenous ADAR expression to achieve robust activation, limiting their translational potential, as ADAR overexpression can result in widespread off-target editing with unknown consequences (*17–21*).

Here, we introduce a new class of genetic switches, termed **adaptamers** (<u>AD</u>AR modulatable <u>aptamers</u>), capable of driving therapeutically relevant transgene expression in vivo. Adaptamers consist of a small-molecule-binding aptamer fused to a short double-stranded RNA region harboring a premature stop codon. Small-molecule binding stabilizes the RNA structure, promoting recruitment of endogenous ADAR, which edits the stop codon and restores translation. These switches are compact (<120 base pairs), tunable, and operate exclusively at the RNA level. While adaptamers function effectively without exogenous ADAR, we further engineered a positive feedback mechanism in which ADAR expression is conditionally induced, enhancing output in tissues with low basal ADAR activity while avoiding constitutive, off-target RNA editing.

Obesity represents a major public health crisis, affecting approximately 40% of adults over the age of 25 in the United States and contributing to severe comorbidities, including type 2 diabetes, cardiovascular disease, and metabolic dysfunction-associated steatohepatitis (*22*, *23*). Fibroblast growth factor 21 (FGF21) has emerged as a promising therapeutic alternative to GLP-1 agonists, owing to its ability to ameliorate obesity and associated metabolic disorders (*24–26*). Unlike GLP-1 agonists, which primarily act by modulating insulin and glucagon secretion and suppressing appetite, FGF21 promotes weight loss by increasing systemic metabolic activity (*26*, *27*). FGF21 gene therapy has also been shown to extend lifespan in mice, largely through metabolic remodeling (*26*). However, FGF21 has a short circulating half-life of approximately two hours, rendering protein-based delivery impractical (*28*). Conversely, constitutive FGF21 overexpression via gene therapy carries significant risks, including muscle atrophy and reduced bone density (*29–31*).

To harness the therapeutic benefits of FGF21 while minimizing these risks, we employed adaptamer-regulated FGF21 expression delivered by AAV. Because edited transcripts persist beyond the presence of the inducing small molecule, this approach enables high-level, stable yet reversible expression in vivo, functionally analogous to a continuous FGF21 infusion. This unique delivery paradigm supports rapid metabolic remodeling in a controlled and safe manner, resulting in robust weight loss and reversal of obesity-associated comorbidities.

## RESULTS

### Engineering small molecule inducible ADAR editing

The adaptamer system was engineered by fusing the cb32 tetracycline minimer—an aptamer with high affinity for tetracycline (*7*, *32*)—to destabilized variants of a naturally occurring RNA structure known to robustly recruit endogenous ADARs (*33*). We hypothesized that tetracycline binding would stabilize the double-stranded RNA region, enabling conditional A-to-I editing of an in-frame stop codon by endogenous ADAR and thereby permitting downstream transgene translation (**Fig. 1a**). Consistent with this model, transient transfection of HEK293T cells with EGFP reporters followed by exposure to 80 μM tetracycline selectively increased expression from adaptamer-containing constructs (**Fig. 1b**). Tetracycline had negligible effects on basal transgene expression, as shown by a no-stop control, or on global ADAR activity, as demonstrated by a no-aptamer reporter (**Fig. 1b**). Drug-induced A-to-I editing at the target stop codon confirmed that activation was editing-dependent (**Fig. 1b**).

**Fig. 1.**
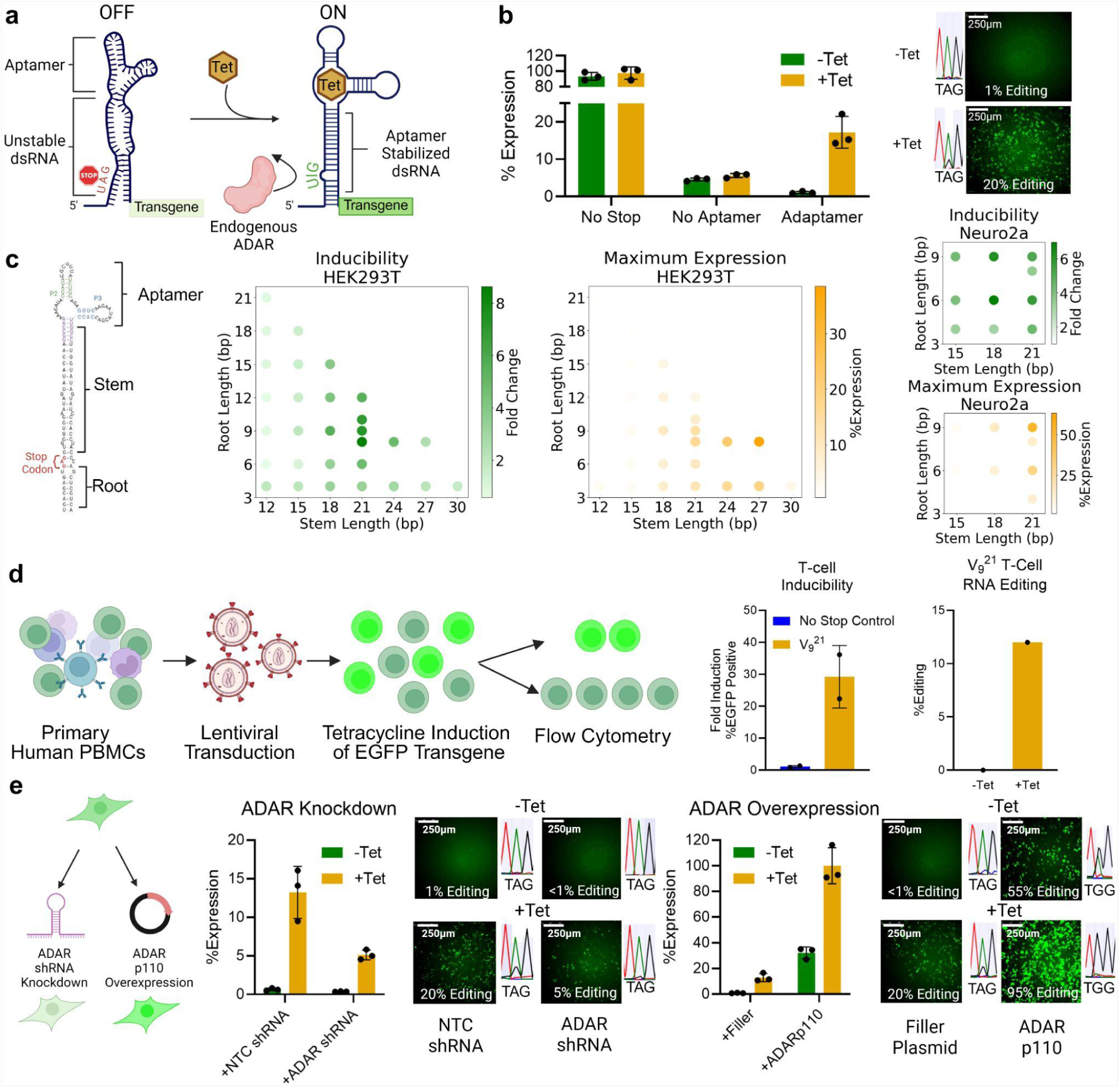
Small-molecule–regulated gene expression via Adaptamers. (a) Schematic of the proposed adaptamer mechanism. Tetracycline binding stabilizes the double-stranded RNA region, enabling recruitment of endogenous ADAR to edit out a stop codon and permit downstream translation. (b) Expression and RNA editing of transgenes in HEK293T cells transfected with the adaptamer construct (V_9_^21^) and cultured with 0 or 80 μM tetracycline, compared to control constructs. (c) Screen of stem and root length variants showing fold change and maximum induced expression in human and mouse cell lines. (d) Transduction of primary human PBMCs (predominantly T cells) with adaptamer-controlled EGFP demonstrates inducible expression. (e) Modulation of cellular ADAR levels via shRNA knockdown or plasmid overexpression impacts adaptamer-driven expression and RNA editing. Error bars represent mean ± s.d. (n = 2–3), and individual dots represent biological replicates. Percent expression is calculated as mean fluorescence intensity normalized to the no-stop-codon control with tetracycline. Maximum expression is percent expression with 80 μM tetracycline. Fold change is maximum expression divided by percent expression at 0 μM tetracycline. Fold induction in T cells was calculated as percent EGFP-positive cells at 80 μM divided by percent at 0 μM tetracycline.

To optimize inducibility and expression, we systematically varied the length of the stem above and the root below the stop codon (**Fig. 1c**). Transient transfection assays in HEK293T cells identified constructs achieving robust tetracycline dependent induction and up to 39% of constitutive expression (**Fig. 1c**). In general, extending the stem and root enhanced maximal expression, consistent with increased thermodynamic stability of longer double-stranded regions (**Fig. 1c**). Based on its favorable balance of maximal expression and fold induction across both HEK293T cells and Neuro2a cells, the V_9_ ^21^ construct (9-base root, 21-base stem) was selected for further study (**Fig. 1c**).

To assess functionality in primary human cells, peripheral blood mononuclear cells–derived T cells were activated, transduced with EGFP-expressing lentiviral vectors, and cultured in the presence or absence of tetracycline prior to flow cytometric analysis (**Fig. 1d**). V_9_ ^21^-transduced T cells exhibited a marked increase in both EGFP-positive cells and RNA editing rates upon tetracycline treatment, confirming robust inducibility in primary human cells (**Fig. 1d**).

Finally, modulation of ADAR abundance in HEK293T cells via shRNA-mediated knockdown or overexpression of ADAR1 p110 revealed a strong correlation between ADAR levels and adaptamer-driven expression (**Fig. 1e**). Importantly, tetracycline remained an effective inducer across all conditions, demonstrating switch functionality over a broad range of endogenous ADAR activity (**Fig. 1e** and **Supplementary Fig. 1)**. RNA editing analysis further showed that tetracycline consistently increased A-to-I editing at the internal stop codon across all ADAR levels, tightly correlating editing efficiency with expression output (**Fig. 1e**).

### Engineering ADAR feedback Loops to Increase Expression

We observed that increasing overall adaptamer length enhanced maximal expression but reduced fold induction due to elevated basal activity (**Fig. 1c**). To overcome this tradeoff, we decided to engineer a positive feedback loop in which adaptamer-regulated exogenous ADAR expression could in turn amplify output while maintaining low background (*34*). This design could also extend adaptamer applicability to settings with limited endogenous ADAR activity, such as immortalized cell lines (*35*). Specifically, the system comprises an adaptamer-controlled transgene followed in series by an adaptamer-controlled ADAR, separated by self-cleaving 2A peptides to ensure independent protein function (**Fig. 2a**).

**Fig. 2.**
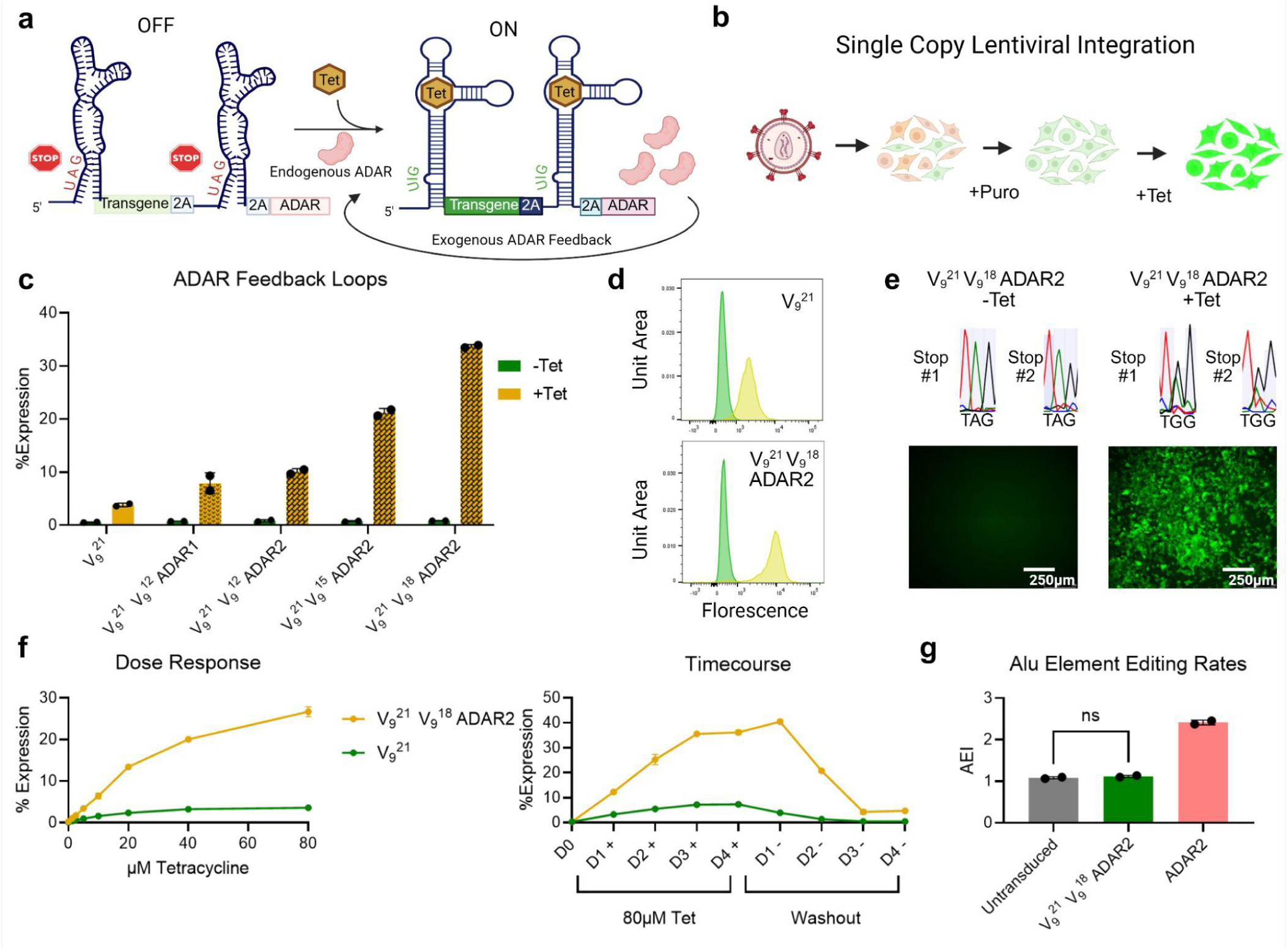
Enhancing adaptamer activity via ADAR feedback loops. (a) Schematic of an adaptamer-controlled transgene followed by a secondary adaptamer regulating exogenous ADAR expression to amplify induced output. (b) Diagram illustrating the generation of single-copy integration feedback loop cell lines to approximate in vivo ADAR expression levels. (c) Expression of feedback loop constructs (black patterning) compared to an adaptamer-only cell line. (d) Representative FACS histograms (log scale) showing induced (yellow) and uninduced (green) feedback loop cell lines versus adaptamer-only cells. (e) RNA editing levels and corresponding fluorescence. (f) Dose-response curves with EC50 values of 17.1 μM for V_9_^21^ and 29.6 μM for V_9_^21^–V_9_^18^–ADAR2, along with time-course data. (g) Global RNA editing rate (AEI) in feedback loop cell lines compared to parental cells and single-copy ADAR2-overexpressing cells (P = 0.32, unpaired two-tailed t-test). Error bars represent mean ± s.d. (n = 2), and individual dots indicate biological replicates. Percent expression is normalized to a single copy constitutive EGFP-overexpressing cell line.

To ensure broad experimental applicability, we generated single-copy lentiviral integration cell lines harboring the engineered adaptamer system (**Fig. 2b**). Initial comparisons revealed that constructs encoding ADAR1 p110 or ADAR2 produced similar protein levels; we therefore selected ADAR2 due to its smaller size (2.2 kb versus 3.7 kb) and reduced off-target editing profile (**Fig. 2c**) (*21*). Using V_9_ ^21^ as the primary regulatory element, we inserted secondary adaptamers (V_9_ ^12^, V_9_ ^15^, or V_9_ ^18^) between the EGFP transgene and ADAR2 to tune effector production while maximizing reporter output. Notably, these feedback-loop cell lines exhibited 2-to 9-fold higher expression and approximately threefold increased RNA editing upon tetracycline induction, without increased basal expression, relative to V_9_ ^21^ alone (**Fig. 2c–e**). Additionally, induction was uniform across the cell population, with no evidence of sporadic background activation (**Fig. 2d,e**). The top-performing construct, V_9_ ^21^–V_9_ ^18^–ADAR2, achieved higher expression than the parental V_9_ ^21^ across all tested tetracycline concentrations, with only a modest increase in EC50 (29.6 μM versus 17.1 μM), indicating that activation remained sensitive and tunable (**Fig. 2f**). Importantly, expression was rapidly reversible within three days following tetracycline withdrawal, exhibiting only a brief delay relative to the parent construct (**Fig. 2f**). Transcriptome-wide RNA-seq analysis revealed no increase in global A-to-I editing rates or differentially edited sites, demonstrating that background exogenous ADAR expression remained sufficiently low to have negligible off-target effects (**Fig. 2e**).

### In vivo gene control via Adaptamers

To evaluate the adaptamer’s capacity for enabling tunable gene therapy in vivo, we generated AAVs expressing adaptamer-controlled firefly luciferase (fLuc). We omitted the ADAR feedback loop due to packaging constraints and the generally higher endogenous ADAR activity in vivo, which we anticipated would enhance maximal expression (*35*). Mice were systemically injected with 1×10¹² vg of AAV9 expressing either V_9_^21^-controlled or constitutive fLuc to assess expression and inducibility. Initial induction cycles with 60 mg/kg tetracycline via intraperitoneal injection—a standard murine dose—demonstrated reversible and repeatable expression increases, returning to near baseline within 48 hours (**Supplemental Fig. 2a,b**) (*7*, *9*).

Six weeks post-injection, mice received 60 mg/kg tetracycline and were imaged at 0, 4, 8, 12, and 24 hours (**Fig. 3a**). To confirm RNA editing at harvest, mice were redosed at 24 hours and sacrificed at 28 hours. Induced expression was detectable within 4 hours, peaking at 12 hours (**Fig. 3b**). Liver lysate analysis of luminescence and RNA editing corroborated IVIS findings (**Supplemental Fig. 2c,d**). Mice transduced with an ADAR-editable loop lacking the tetracycline aptamer showed no increase in RNA editing, confirming adaptamer-specific induction (**Supplemental Fig. 2e**).

**Fig. 3.**
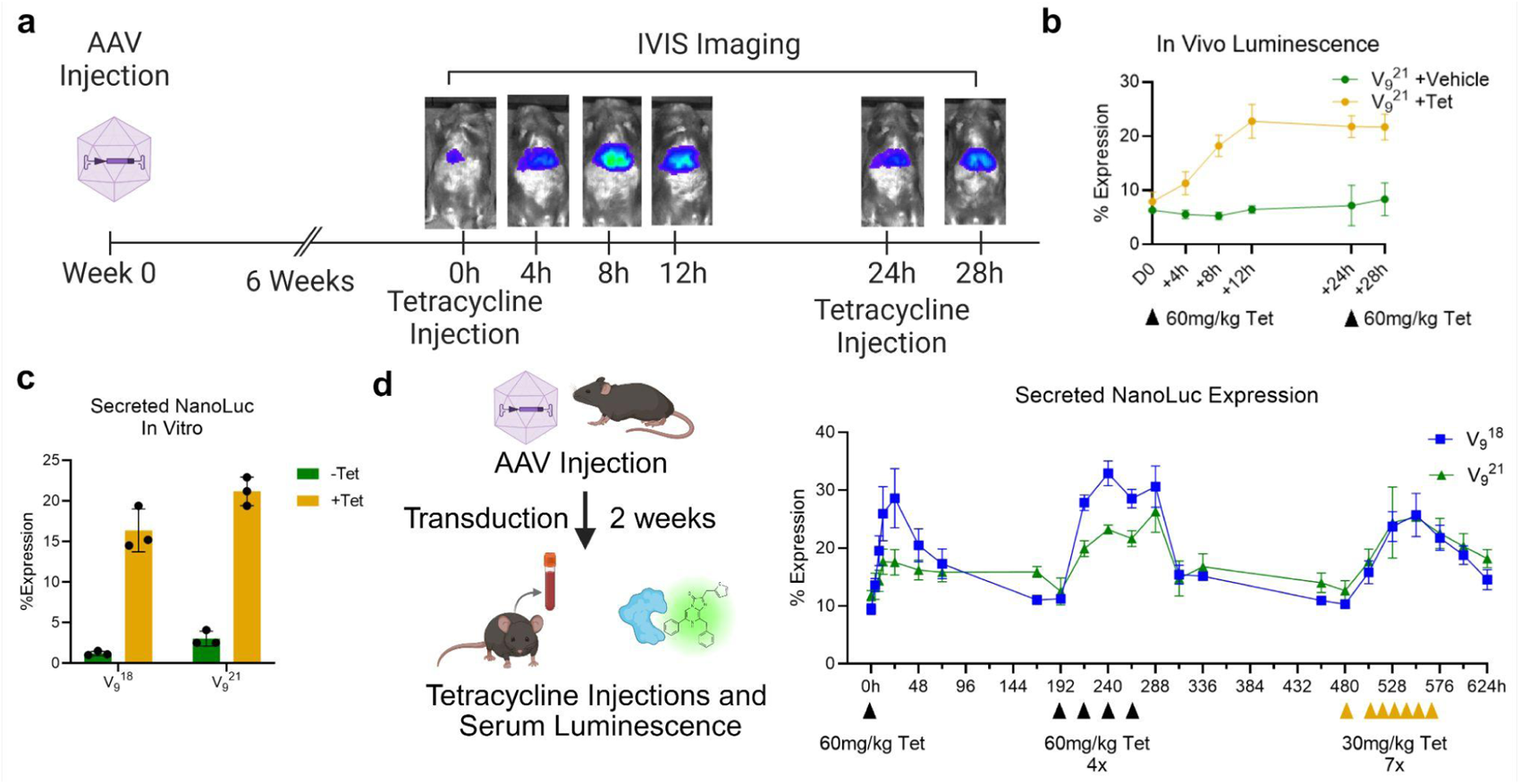
In vivo adaptamer-mediated transgene expression. (a) Schematic of inducible firefly luciferase expression in vivo. Mice were injected with adaptamer-controlled AAV9. After transduction equilibration, mice received 60 mg/kg tetracycline, were imaged at defined intervals, and then received a second tetracycline dose at 24 hours, 4 hours prior to tissue harvest. (b) Luminescence over time following tetracycline administration. (c) In vitro secreted NanoLuc expression in HEK293T cells comparing V_9_^18^ and V_9_^21^ constructs. (d) In vivo kinetics of secreted NanoLuc for V_9_^18^ and V_9_^21^ across multiple tetracycline induction regimens. Error bars represent mean ± s.e.m. (n = 3), and individual dots indicate biological replicates. Percent expression is normalized to the no-adaptamer control also treated with tetracycline.

To rapidly optimize adaptamer designs for improved in vivo functionality, we next screened multiple designs by injecting a pool of adaptamer variant-containing AAVs. We dosed half the animals with tetracycline, and quantified differential RNA editing rates using each adaptamer sequence as its own barcode (**Supplemental Fig. 2f**). From this screen we selected V_9_ ^18^, which has a three-base shorter stem than V_9_ ^21^ and reduced background in vitro as measured with a secreted NanoLuc (sNLuc) reporter (**Fig. 3c**). Using AAV8 expressing inducible sNLuc, we compared V_9_ ^18^ and V_9_ ^21^ in vivo, confirming lower background and similar or improved induced expression across multiple tetracycline dosing regimens for the V_9_ ^18^ design (**Fig. 3c**). Secreted NanoLuc expression peaked at 24 hours—slightly later than fLuc, likely due to secretion and biodistribution—allowing repeated tetracycline dosing to achieve sustained induced expression (**Fig. 3c**).

### Adaptamer mediated inducible FGF21 gene therapy for reversing obesity

To demonstrate therapeutic potential, we next applied the adaptamer system to inducibly overexpress FGF21 for treatment of chronic obesity. FGF21 overexpression is known to enhance adipose metabolism, promote liver rejuvenation, and improve glucose tolerance but can dysregulate muscle and bone metabolism, highlighting the need for inducible control (*29–31*). We first optimized viral dosing by injecting mice with 5×10¹⁰, 2×10¹¹, or 1×10¹² vg of inducible AAV8-FGF21, selecting the midpoint based on prior AAV studies and our observed ∼10% background from sNLuc experiments (**Supplementary Fig. 3a**) (*26*, *36*). The 2×10¹¹ vg dose yielded negligible basal FGF21 yet achieved therapeutically relevant levels upon 60 mg/kg intraperitoneal tetracycline administration (**Supplementary Fig. 3a**). Next, we tested three tetracycline regimens—one 60 mg/kg injection, two 30 mg/kg injections 12 hours apart, and a single 30 mg/kg injection with six days between doses—to determine minimal effective dosing. All regimens produced comparable FGF21 expression 24 hours post-induction, so we proceeded with the conservative 30 mg/kg dose (**Supplementary Fig. 3b**).

To model obesity therapy, 21-week-old diet-induced obese (DIO) mice were injected with 2×10¹¹ vg of inducible AAV8-FGF21 and treated one week later with daily 30 mg/kg tetracycline to trigger metabolic reprogramming (**Fig. 4a**). Inducible FGF21 expression achieved sustained therapeutic levels in drugged mice while remaining low in vehicle-treated controls (**Fig. 4b**). Induction persisted for at least 24 hours post-tetracycline, with elevated FGF21 and RNA editing despite FGF21’s short half-life (*28*) and rapid tetracycline clearance (*7*)(**Supplementary Fig. 4a**). Substantial weight loss occurred only in mice receiving both AAV-FGF21 and tetracycline (FGF21 +Tet), confirming the requirement for both gene therapy and induction (**Fig. 4c, Supplementary Fig. 4b**). Glucose tolerance testing on Day 22 further confirmed improved glycemic control in FGF21 +Tet mice, reflecting effective metabolic reprogramming (**Fig. 4d**). Histological analysis showed near-complete reversal of hepatic steatosis, evidenced by elimination of lipid droplets in hepatocytes (**Fig. 4e**). Additionally, lipid accumulation in interscapular brown adipose tissue decreased, and inguinal white adipose tissue exhibited reduced adipocyte size (**Supplementary Fig. 4b,d**).

**Fig. 4.**
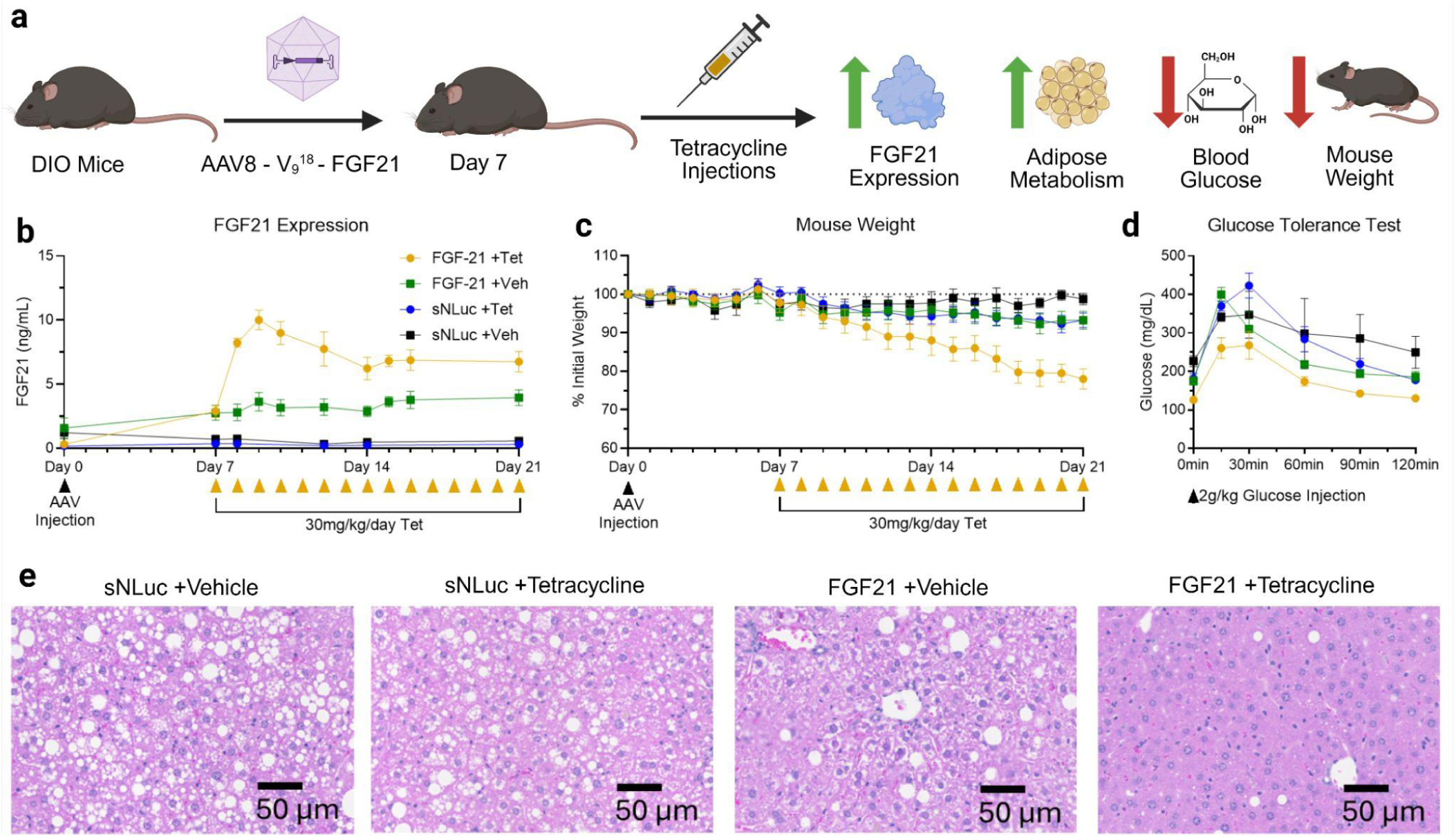
Adaptamer-mediated inducible FGF21 gene therapy reverses obesity. (a) Schematic of the FGF21 gene therapy strategy, where inducible FGF21 expression enhances fat metabolism, improves glucose tolerance, and reduces body weight in obese mice. (b) Serum FGF21 levels in diet-induced obese (DIO) C57BL/6 mice following injection of 2×10¹¹ vg per animal, measured by ELISA. (c) Changes in body weight expressed as a proportion of initial weight at the time of AAV injection. (d) Glucose tolerance test performed on Day 22. (e) Hematoxylin–eosin staining of liver sections showing reversal of hepatic steatosis in FGF21 + tetracycline–treated mice. Error bars represent mean ± s.e.m. (n = 4).

## DISCUSSION

While gene therapy offers the potential for long-lasting treatment, current modalities suffer from a “Goldilocks problem”—constitutive expression can lead to toxicity (e.g., FGF21-induced bone loss or muscle atrophy) or desensitization, while transient vectors fail to provide durable relief. To address this, we introduced the **Adaptamer**, a compact (<120 bp), human-compatible, small-molecule–responsive riboswitch that enables inducible and reversible control of gene expression via endogenous ADAR-mediated RNA editing. Unlike existing systems that rely on immunogenic bacterial proteins (e.g., Tet-On) or RNA cleavage, the Adaptamer stabilizes a double-stranded RNA structure upon ligand binding to recruit ADAR, converting a premature stop codon into a sense codon and restoring translation. This system differs from other riboswitches in three key ways, providing unique advantages for both utility and future development. First, it is the first riboswitch, to our knowledge, to use a non-cleavage, non-splicing mechanism, enabling RNA-based delivery and extra-nuclear activation. This allows integration with mRNA lipid nanoparticles and lentiviral vectors, expanding delivery options and enabling cell-based screens. Second, the core adaptamer is extremely compact, facilitating efficient integration into space-constrained vectors such as AAVs carrying large transgenes, including full-length saCas9. Unlike other riboswitches with large, GC-rich structures, particularly splicing-based designs, the adaptamer is transgene-agnostic and streamlined for vector construction (*9*). Finally, the system is modular, with distinct effector and small-molecule binding domains, which we anticipate will allow substitution of the tetracycline aptamer with other ligand-binding aptamers without major scaffold modifications. Expression is tunable via root and stem truncation or extension, enabling precise control of dynamic range. Leveraging A-to-I editing as the effector enabled the first reported in vivo pooled riboswitch screen, a strategy for rapid testing across tissues and contexts.

The adaptamer also advances ADAR-mediated sensing. While previous ADAR-based switches respond to endogenous molecules, this is the first system controlled by an FDA-approved exogenous ligand and capable of producing therapeutically relevant transgene levels without exogenous ADAR overexpression (*17*, *18*, *20*). Its reliance on endogenous ADAR offers additional advantages: higher brain ADAR activity may allow tissue-specific expression and liver detargeting (*14*), while inducible ADAR p150 expression in immune contexts, such as activated T cells, could enable synergistic protein production in combination with a small molecule (*37*, *38*).

Crucially, the adaptamer demonstrates clinical relevance. In a diet-induced obesity mouse model, small-molecule-induced FGF21 expression produced therapeutic benefits without background activity, reversing obesity, glucose intolerance, and hepatic steatosis. Unlike cleavage-based RNA switches, adaptamer RNA remains stable without the small molecule, permitting sustained yet inducible expression—critical when short inducer exposure windows are required, as with tetracycline (*7*). This inducible stability enhances both the safety and efficacy of FGF21 gene therapy.

Despite these advances, limitations remain. Dynamic range and maximal expression can be improved through mutational screens and directed evolution, enabled by the cleavage-free design. Additionally, the current system is limited to tetracycline, restricting multiplexing. As new aptamers are identified via SELEX, they can be incorporated to broaden adaptamer versatility with minimal scaffold modifications (*8*, *39*).

In conclusion, we demonstrate that adaptamers can be rationally engineered to enhance functionality, and deployed in vivo to deliver therapeutic benefits e.g. through small-molecule-controlled metabolic reprogramming. While further optimization will enhance performance, the adaptamer system represents a novel advance in synthetic gene regulation, expanding the capabilities of both riboswitches and ADAR-mediated sensors.

## ACKNOWLEDGEMENTS

We thank members of the Mali lab for discussions, advice, and help with experiments. We thank Sami Nourreddine for useful discussions and support throughout all stages of the study. We thank Kristen Jepsen, Director of IGM Genomics Center UC San Diego. We thank Zbigniew Mikulski, Director of The La Jolla Institute for Immunology Microscopy & Histology Core. This work was generously supported by NIH grants (R01HG012351, R01NS131560), a Department of Defense Grant (W81XWH-22-1-0401), and UCSD Institutional Funds. Some schematics were created using BioRender.

## DECLARATION OF INTERESTS

The authors have filed a patent based on this work. P.M. is a scientific co-founder of Shape Therapeutics, Boundless Biosciences, Navega Therapeutics, Pi Bio, and Engine Biosciences. The terms of these arrangements have been reviewed and approved by the University of California San Diego in accordance with its conflict of interest policies.

## DATA AVAILABILITY

All key reagents will be made available via Addgene. Source data are provided with this paper.

## METHODS

### Plasmid Design/Construction

Parental plasmids were constructed by overlap extension PCR of relevant components and Gibson assembly into either pX601-AAV-CMV (Addgene #61591), ePIP (Addgene #172110), or a custom synthesized rAAV2 vector backbone. Adaptamer sequences were chemically synthesized as eBlocks by IDT and inserted into backbones through Gibson assembly.

### Cell Culture

HEK293T and HeLa cells were cultured in DMEM (Thermo Fisher Scientific) supplemented with 10% FBS (Thermo Fisher Scientific) and 1% Antibiotic-Antimycotic (Thermo Fisher Scientific) in an incubator at 37°C and 5% CO_2_ atmosphere.

Neuro2a cells were cultured in EMEM (ATCC) supplemented with 10% FBS (Thermo Fisher Scientific) and 1% Antibiotic-Antimycotic (Thermo Fisher Scientific) in an incubator at 37°C and 5% CO_2_ atmosphere.

### General Transfection Protocol

24 hours prior to transfection cells were seeded at 25,000 cells per square cm in either a 6, 24, or 48 well plate. The following day cells were transfected with 250ng plasmid and 0.5µL Lipofectamine 2000 (Thermo Fisher Scientific) per cm^2^ according to manufacturers protocol. Media was exchanged 4 hours after transfection with the appropriate tetracycline concentration.

### Flow cytometry

Harvest for flow cytometry occurred 24 hours after transfection for HEK293T cells, 48 hours after transfection for N2a cells, or at varying timepoints for transduced HEK293T cells. Growth media was removed and 50µL per cm^2^ 0.05% Trypsin (Thermo Fisher Scientific) was added for a 5 minute incubation at 37C to dissociate. Cells were resuspended in a solution of 50% culture media and 50% PBS (Thermo Fisher Scientific) to a final concentration of approximately 4e5 cells per mL. 10,000 single cells per sample were analysed with a BD FACSCanto or LSRFortessa machine. Using untreated cells as a gating control, live cells were gated using forward and side scatter, single cells were gated using both side and forward scatter area versus width. Percent expression was calculated using the Mean Fluorescence Intensity of transfected cells or Median Fluorescence Intensity of Transduced cells, subtracting untreated cell value as background and normalizing to no adaptamer control MFI.

### RNA editing analysis

Cells were lysed 48 hours after transfection and processed using the RNAeasy Mini Kit (Qaigen #74104) with optional on column DNAse digestion following manufacturer protocol. RNA was extracted from manually homogenized mouse tissue using TRIzol (Invitrogen) and chloroform according to manufacturer protocol and process using the RNAeasy Mini Kit. cDNA was generated using the Protoscript® II First Strand cDNA Synthesis Kit (NEB E6560L). The adaptamer sequence was amplified using primers 3xFlag_F: 5’GACTACAAAGACCACGACGGGGAC3’ and P2A_R: 5’CGGTCCAGGATTCTCTTCGACATC3’ or Adapt_Univ_R: 5’CTCGACCAGGATGGGCAC3’ and Sanger sequenced using the appropriate reverse primer.

### ADAR Modulation

12,500 HEK293T cells per well were seeded in a 48 well plate 24 hours prior to transfection with 250ng of ADAR1 shRNA, ADAR p110, or pUC19 plasmid using 0.25µL of Lipofectamine 2000 (Thermofisher) per well. 16 hours after transfection with ADAR modulators, a gentle media exchange was performed followed by a transfection of 250ng of EGFP expressing adaptamer constructs using 0.25µL of Lipofectamine 2000 (Thermofisher) per well. 4 hours after EGFP plasmid transfection a complete media exchange was performed with media containing either 0 or 80µM tetracycline. Flow cytometry was performed at 24 hours post media change and RNA was harvested for editing analysis at 48 hours. ADAR transcript qPCR was performed using a CFX Connect Real Time PCR Detection System (Bio-Rad) and iTaq Universal SYBR Green Supermix (Bio-rad) with primers hADAR1-qPCR-F: 5’TCCGTCTCCTGTCCAAAGAAGG3’ and hADAR1-qPCR-R 5’TTCTTGCTGGGAGCACTCACAC3’ and normalized to B-actin expression with primers qPCR_human_B-ACTIN_F: 5’CATGTACGTTGCTATCCAGGC3’ and qPCR_human_B-ACTIN_R: 5’CTCCTTAATGTCACGCACGAT3’.

### Primary Human PBMC Transduction

To produce concentrated lentivirus an 80% confluent 15cm plate of HEK293T cells with fresh media was transfected with 3μg pMD2.G, 12μg of pCMV delta R8.2, and 9μg of EGFP expressing lentiviral vector using 36μL of Lipofectamine 2000 (ThermoFisher Scientific). 48 hours after transfection media was collected at 4C and replaced. At 72 hours the second batch of media was collected, pooled with the first batch, and centrifuged at 500xg for 5 minutes to remove carryover cells. The supernatant was then passed through a 0.45μm filter and concentrated in a 100 kDa MWCO Ultricentrifugal Filter (MilliporeSigma) to a final volume of ∼400μL. Human PBMCs (Stemcell Technologies) were thawed three days prior to transduction and seeded at 1e6 cells/mL in RPMI 1640 (Thermo Fisher Scientific) supplemented with 10% FBS (Thermo Fisher Scientific), 1% Antibiotic-Antimycotic (Thermo Fisher Scientific), 200 IU/mL rhIL-2 (Stemcell Technologies) and ImmunoCult™ Human CD3/CD28 T Cell Activator (Stemcell Technologies). Cells were centrifuged and reseeded in fresh media 24 hours prior to transduction. 50μL of concentrated virus was mixed with 1e6 cells and polybrene added to 8μg/mL(MilliporeSigma). Cell virus mixture was spinnoculated at 1000xg and 32C for 90 minutes, then supernatant was removed and cell pellet was resuspended and seeded in RPMI 1640 (Thermo Fisher Scientific) supplemented with 10% FBS (Thermo Fisher Scientific), 1% Antibiotic-Antimycotic (Thermo Fisher Scientific), 200 IU/mL rhIL-2 (Stemcell Technologies) at 2e6 cells per mL. Two days after transduction, PBMCs were split, centrifuged, and resuspended in complete RPMI without activator containing either 0μM or 20μM tetracycline. Two days after addition of tetracycline, cells were analysed via flow cytometry and percent EGFP positive cells were gated using an untransduced PBMC sample as a negative control. RNA was also extracted from cells and used to determine RNA editing rates.

### Lentiviral HEK293T Cell Line Generation

For each lentiviral line, one well of a 6 well plate was seeded with 100,000 HEK293T cells 24 hours prior to Lipofectamine 2000 transfection with 200ng pMD2.G, 800ng of pCMV delta R8.2, and 600ng of lentiviral vector with a media exchange immediately prior to transfection. 24 hours after transfection media was collected, mixed with polybrene (MilliporeSigma) to 8 μg/mL, centrifuged at 500xg for 5 minutes to remove cells, and used to replace media on a new well of HEK293T cells seeded one day prior. The following day lentivirus containing media was replaced with standard complete media, and two days after transfection cells were cultured in media containing 1 μg/mL puromycin (ThermoFisher Scientific) for the remainder of experiments to select for transduced cells.

### RNAseq and global ADAR editing analysis

A confluent well of a 6 well plate of transduced cell lines was collected and RNA extracted using the RNAeasy Mini Kit (Qaigen #74104). A NEBNext Poly(A) mRNA Magnetic Isolation Module Kit (NEB E7490S) was used to select from 1 μg per sample of total RNA and Illumina-compatible RNAseq libraries were created using an NEBNext Ultra RNA Library Prep Kit (NEB E7530S). Sequencing was performed on an Illumina NovaSeq X, with paired end 100 base pair reads. Read files were aligned with BWA-MEM2 to the hg38 Human Reference genome. For RNA editing analyses, Alu Editing Index (AEI) was determined using RNAEditingIndexer with default parameters and the builtin hg38 human genome reference, alu annotations, and SNPs(*40*). Differentially edited sites were determined via LoDEI using default parameters (q < 0.1) and hg38 Human Genome Annotation (2024-A) with GENCODEv44 annotations(*41*).

### AAV Production

AAV virus was produced by triple-transfection of HEK293T cells and purified with an iodixanol gradient as previously described(*42*, *43*). Briefly, cells were seeded at 10% confluency in 15cm dishes 2 days prior to transfection, with complete media exchanges performed 2 hours before and 16 hours after transfection. Each plate was transfected with 30 μg total of transgene vector, capsid vector (pXR-9 or pXR-8), and pHelper vector in an equimass ratio utilizing linear polyethylenimine (PEI) at a ratio of 4:1 PEI to DNA (PEI dissolved 1 mg/mL in PBS, pH balanced to 7 with HCl and NaOH). 4 days post-transfection virus was harvested from supernatant via overnight 4C 10% polyethylene glycol (PEG) incubation and directly from freeze-thaw lysed cells. The combined virus was Benzonase (SigmaAldrich) treated at 37 C for 1hr to digest any unencapsulated and an iodixanol gradient was used to isolate virus from the supernatant. Virus was then dialyzed using 50 kDa MWCO centrifugal filters (MilliporeSigma) in a solution of PBS (ThermoFisher Scientific), 50 mM NaCl, and 0.0001% of Pluronic F68 (ThermoFisher), concentrated to a final volume of ∼200 μL, and quantified via qPCR compared to a known standard (ATCC VR-1616) utilizing serial dilutions and primers targeting the ITR regions, AAV-ITR-F: 5’-CGGCCTCAGTGAGCGA-3’ and AAV-ITR-R: 5’-GGAACCCCTAGTGATGGAGTT-3’.

### Animal Studies

All animal procedures were performed in accordance with protocols approved by the Institutional Animal Care and Use Committee of the University of California, San Diego. All mice were acquired from Jackson Labs. Adult male C57BL/6J mice were injected retro-orbitally with 1e12vg of AAV virus unless otherwise noted.

For blood collection and serum extraction mice were anesthetized using controlled O_2_/isoflurane and ∼20µl of blood was collected via a tail nick. Blood was allowed to coagulate for 15 minutes at room temperature then was centrifuged at 2500x g for 10 minutes to separate serum. Serum was removed and frozen at -80C for processing at a later time.

### Tetracycline Injections

Tetracycline injection solutions were prepared immediately before injection as previously described(*7*). Briefly, 12 mg of tetracycline hydrochloride (SigmaAldrich) was dissolved by vortexing in 0.8mL of a 25% weight by volume solution of B-cyclodextran (MilliporeSigma) in PBS, then diluted by addition of 1.2mL of PBS. Mice were injected intraperitoneally, alternating injection sites to minimize discomfort.

### IVIS Imaging

Mice were anesthetized using controlled O_2_/isoflurane and injected with 10 µL of a 15 mg/ml solution of d-Luciferin (GoldBio, LUCK) in PBS per gram body weight. 5 minutes after injection mice were imaged every minute with 0.5 second exposure and 4x4 binning using a Perkin Elmer IVIS Spectrum system until 20 minutes post injection. For quantification consistency the liver was gated and the maximum observed average radiance (photons per second per cm^2^) was taken for each animal at each timepoint. A mouse not injected with AAV was subtracted as background and percent expression was calculated at each timepoint by dividing by the average radiance across three mice injected with AAV9 expressing constitutive firefly luciferase.

### Secreted NanoLuciferase Assay

2µL of serum was diluted 1:50 in PBS, then 10µL of diluted serum was mixed with 10µL of complete substrate from the Nano-Glo® Luciferase Assay System (Promega) and read on a Spectamax iD3 plate reader (Molecular Devices). Serum from a mouse not injected with AAV was subtracted as background and percent expression was calculated at each timepoint by dividing by the average luminescence value of three mice injected with AAV8 expressing constitutive secretedNanoLuc.

### In vivo RNA editing Screen

Six mice were injected with 1e12vg each of a pooled library of AAV encapsulated adaptamers. Two weeks later three animals were injected with two doses of 60mg/kg tetracycline 24 hours apart. All mice were harvested 4 hours after the last dose. Targeted RNAseq of adaptamer library in the liver was used to determine A to I editing rates of each construct in each mouse. Amplicons were amplified from cDNA and processed via Amplicon-EZ (Azenta Life Sciences). Primers, Adapt-Screen-F: 5’ACACTCTTTCCCTACACGACGCTCTTCCGATCTANGAATTCGCCACCATGGACTACAAAG3’, with the N nucleotide varied for multiplexing, and Adapt-Screen-R: 5’GACTGGAGTTCAGACGTGTGCTCTTCCGATCTCAGGCTCGACCAGGATGGGCAC3’. The ratio of reads with an A to I edit to the total number of reads for each construct was calculated to determine RNA editing rate.

### FGF21 Gene Therapy

21 week old DIO C57BL/6J mice were purchased from Jackson Labs and injected with 2e11vg of AAV8 virus after one week of acclimatization. Mice were fed ad libitum with a high fat diet (60% calories from Fat, Bioserv F3282). Animals were placed in weight matched groups on the day of viral injection and cages contained mixed groups to eliminate cage based effects.

### FGF21 ELISA

Serum FGF21 was quantified using Mouse FGF21 DuoSet ELISA kit (R&D DY3057) and an ELISA Starter Kit (Bethyl Laboratories E101) according to manufacturer protocols. Briefly, a clear bottom 96 well ELISA plate was coated overnight with 100µL of 4 µg/mL Capture Antibody diluted in PBS per well, washed 3 times with wash buffer (50 mM Tris buffered saline, pH 8.0, 0.05% Tween20; Sigma Chemical # T9039), then incubated for two hours with 300µL Blocking Buffer (50 mM Tris buffered saline, pH 8.0, 1% BSA; Sigma Chemical # T6789). Plates were washed again three times then 100µL of sample (serum 1:100 dilution) or standard diluted in Sample Diluent (Blocking buffer +0.05% Tween20) and incubated at room temperature for two hours. Plates were washed as previously described then incubated for two hours with 100µL per well of 100ng/mL Detection Antibody in Sample Diluent. Plates were incubated with for 20 minutes in the dark with 100µL of Streptavin-HRP after another set of washes, then incubated for 20 minutes in the dark with 100µL of TMB One Component HRP Microwell Substrate after a final set of washes. Reactions were stopped using 0.16M sulfuric acid stop solution and absorbance reading taken at 450nm with a 570nm correction using a Spectamax iD3 plate reader (Molecular Devices). Values were interpolated using a four-parameter logistic curve fitted to the standard values, averaging technical duplicates.

### Glucose Tolerance Test

Mice were fasted 8 hours overnight prior to testing. Blood and serum were collected immediately before intraperitoneal injection of 2g/kg dextrose (Sigma-Aldrich) dissolved in PBS. For subsequent timepoints only a small drop of blood was collected. Glucose measurements were performed using a handheld glucose monitor (Metene TD-4116).

### DIO Mouse Histology

DIO mice were sacrificed 24 hours after last tetracycline dose and liver, interscapular brown adipose tissue (iBAT), and inguinal white adipose tissue (iWAT) were collected, briefly washed in ice cold PBS, then fixed in Neutral Buffered Formalin, 10% (SigmaAldrich) for 24 hours. After fixation samples were Paraffin-Embedded, sectioned, hematoxylin–eosin stained, and imaged at 20x magnification by the Microscopy and Histology Core of the La Jolla Institute of Immunology.

---

**Supplementary Fig. 1.**
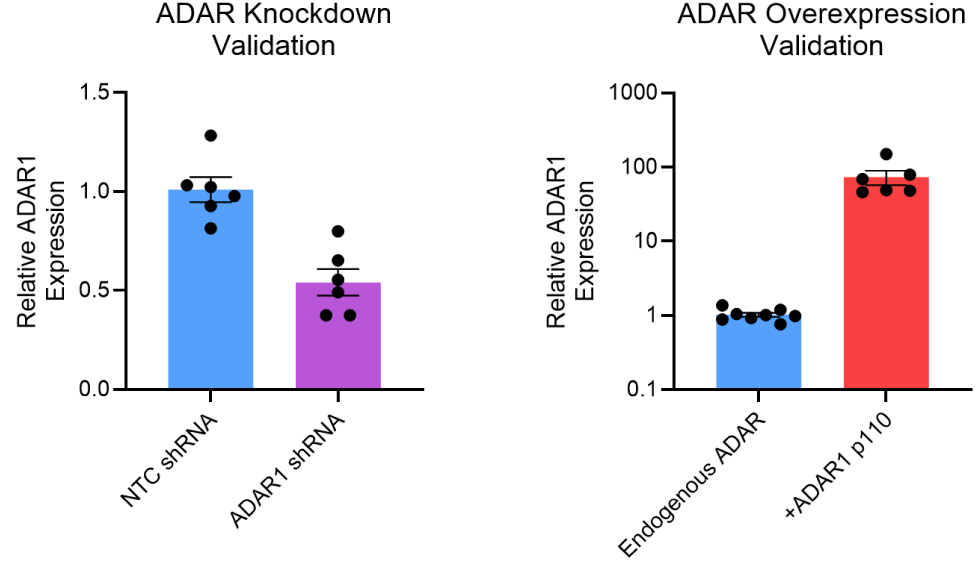
Validation of ADAR modulation. Transfection of ADAR1-targeting shRNA reduced endogenous ADAR transcript levels by ∼50%, whereas transfection with an ADAR1 p110–expressing plasmid increased transcript levels by several orders of magnitude. Error bars represent mean ± s.e.m., and individual dots indicate biological replicates. Relative expression is normalized to the average of non-targeting control (NTC) shRNA or endogenous ADAR conditions.

**Supplementary Fig. 2.**
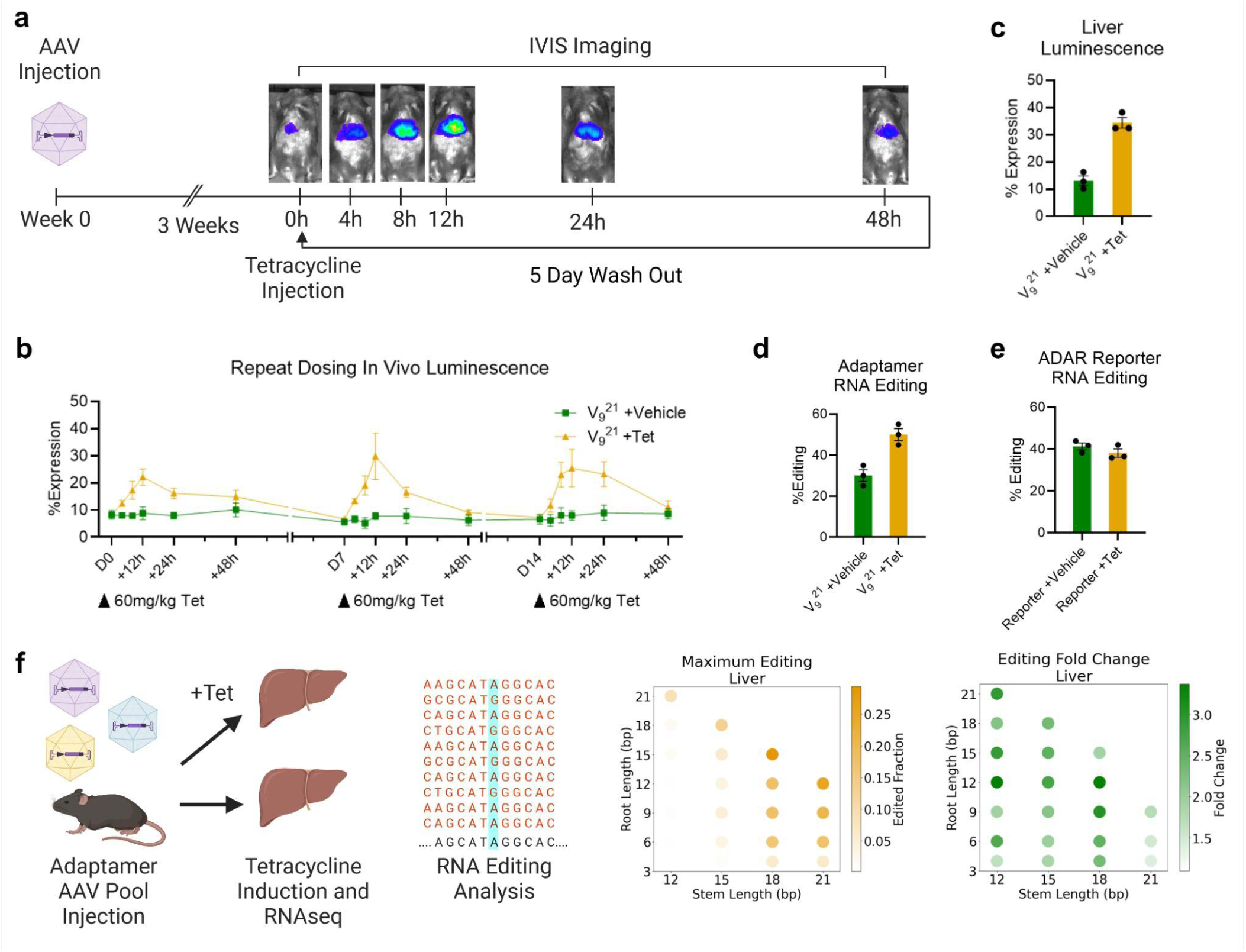
Extended data on adaptamer-mediated inducible in vivo firefly luciferase. (a) Schematic of repeat induction experiment in vivo. Mice were dosed with 60 mg/kg tetracycline, bioluminescence was imaged at defined intervals, followed by a five-day washout and repeat dosing. (b) In vivo luciferase luminescence following multiple tetracycline doses. (c) Luminescence from lysed liver tissue, demonstrating protein-level inducibility. (d) Differential RNA editing rates of the adaptamer at harvest. (e) Effect of intraperitoneal tetracycline on A-to-I editing in the liver of mice transduced with an ADAR reporter construct lacking the tetracycline-binding aptamer. (f) Mice injected with a pooled adaptamer AAV library received either tetracycline or saline, and liver RNA-seq was used to quantify maximum RNA editing and tetracycline-induced increases. Error bars represent mean ± s.e.m. (n = 3), and individual dots indicate biological replicates. Percent expression is normalized to the no-stop-codon control also treated with tetracycline.

**Supplementary Fig. 3.**
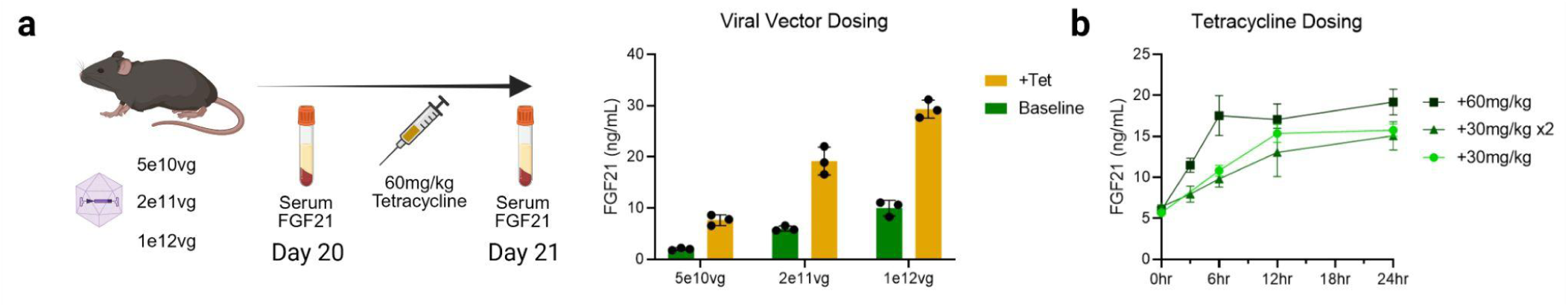
Adaptamer-mediated FGF21 gene therapy dosing optimization. (a) C57BL/6J mice were injected on Day 0 with three different titers of AAV-V918-FGF21 and received 60 mg/kg tetracycline on Day 20 to induce expression. Serum FGF21 levels quantified by ELISA show inducible expression across all viral doses, with variable magnitudes. (b) Three tetracycline dosing regimens produced distinct temporal expression profiles but resulted in similar FGF21 levels at 24 hours post-induction. Error bars represent mean ± s.e.m. (n = 3), and individual dots indicate biological replicates.

**Supplementary Fig. 4.**
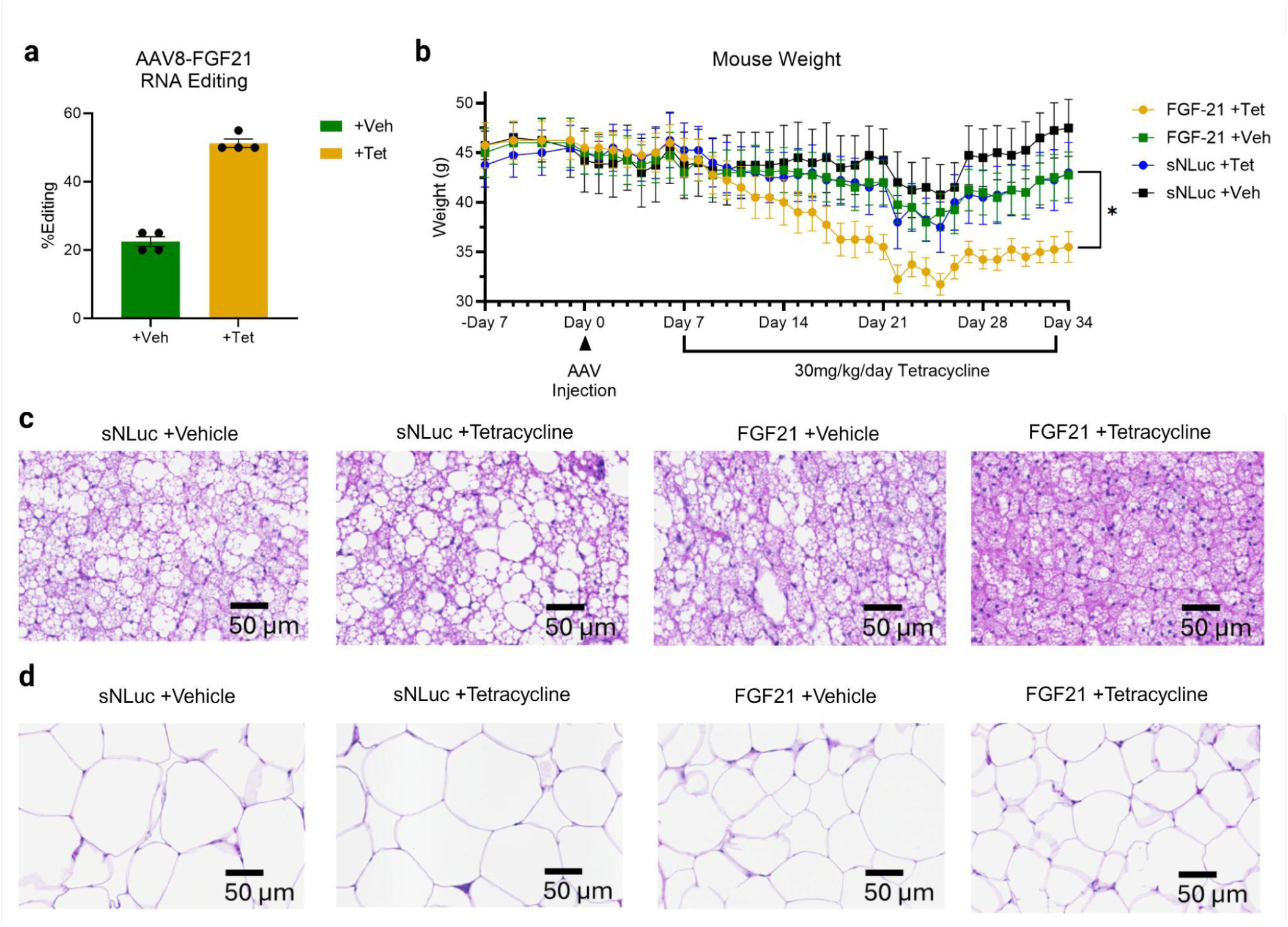
Extended data on adaptamer-mediated inducible FGF21 gene therapy in DIO mice. (a) RNA editing rates measured 24 hours after the final tetracycline dose. (b) Raw body weights of mice over the full experimental course. (c) Hematoxylin–eosin staining of interscapular brown adipose tissue (iBAT) showing reduced lipid deposits in FGF21 + tetracycline–treated mice. (d) Hematoxylin–eosin staining of inguinal white adipose tissue (iWAT) showing decreased adipocyte size in FGF21 + tetracycline–treated mice. Error bars represent mean ± s.e.m. (n = 4), and individual dots indicate biological replicates. *P < 0.05, unpaired two-tailed t-test.

